# Standardizing marine habitat modelling practices to enhance inter-comparability across biological observations

**DOI:** 10.1101/2024.09.02.610745

**Authors:** Alexandre Schickele, Corentin Clerc, Fabio Benedetti, Daniele De Angelis, Urs Hofmann-Elizondo, Matthias Münnich, Jean-Olivier Irisson, Meike Vogt

## Abstract

In recent years, the volume of accessible marine pelagic observations has increased exponentially and now incorporates a wealth of new data types, including information derived from metagenomics and quantitative imaging. This calls for standardized modelling protocol across taxonomically harmonized observations, to better predict biogeographic patterns in space and time, and thus investigate marine ecosystem structure and functioning on a macroecological scale. In this context, we introduce CEPHALOPOD (Comprehensive Ensemble Pipeline for Habitat modelling Across Large-scale Ocean Pelagic Observation Datasets), a standardized and flexible framework to perform multi-species marine habitat modelling across data types and data sources. We built this new framework on observational data from federating initiatives such as AtlantECO, OBIS, GBIF, associated with already existing statistical and machine learning methods that enable to extract and model information from heterogeneous, scarce, and biased field observations. Here, we first document our statistical ensemble modelling approach and then assess its strength and limitations with a virtual ecologist approach. We show how our framework performs in reproducing a range of distributions from biased field samples. Then, we illustrate its performance and comparability across data types by investigating the global diversity patterns of coccolithophores from both abundance and metagenomic data. Our modelling framework serves as a foundation for the consistent generation of Essential Biodiversity and Ocean Variables (EBVs and EOVs) and carries the potential to significantly advance our comprehension of biodiversity and marine ecosystems functioning. Finally, it provides an unprecedented opportunity to foster collaborations in the field of marine science, sustainable ecological practices, and, ultimately, contribute to the preservation of global marine biodiversity.

## INTRODUCTION

Pelagic organisms including plankton and fish, structure marine food webs and sustain essential ecosystem services such as carbon sequestration, oxygen production, fisheries, and associated food security (Cury et al., 2000; Guidi et al., 2016). These organisms are incredibly diverse and their presence or absence, taxonomic or functional composition, abundance and biomass are all accepted indicators of ocean ecosystem structure, health and resilience to climate change, known as EBVs and EOVs (i.e., Essential Ocean and Biodiversity Variables; Muller-Karger et al., 2018). Our historical understanding of plankton and fish biogeography, and the production of associated EBVs and EOVs, relies on decades of observations (Miller et al., 2019). However, marine biodiversity remains critically understudied, with thousands of species whose spatial distribution remains unknown (e.g., Carradec et al., 2018; Peck et al., 2021).

Nowadays, the volume of biological data collected increases faster, thanks to the development of quantitative imaging and genome-based approaches (Delmont et al., 2022; Lombard et al., 2019). This sampling effort largely benefits from the recent development of online data repositories (e.g., Martín Míguez et al., 2019; Mitchell et al., 2020), that help bridge the gap between ecological research and big data science (LaDeau et al., 2017). The development of FAIR principles (i.e., Findable, Accessible, Inter-operable, Reusable; Tanhua et al., 2019) also increased both inter-comparability and inter- operability of such large marine datasets. Nevertheless, their potential to fill knowledge gaps in marine biodiversity and ecosystem functioning remains hampered by their heterogeneity regarding spatio- temporal coverage and resolution, accessibility, metadata harmonization, and specific ecological processes associated with each data type (Dallas & Hastings, 2018; Obiol et al., 2020). Therefore, spatial inference of EOVs and EBVs requires the development of new frameworks that source, combine and integrate this wealth of marine biological datasets across biodiversity metrics, data types, spatio- temporal scales and taxa (Miller et al., 2019).

Proposed decades ago by G.E. Hutchinson (1957), niche theory progressively translated into the development of species distribution models (SDMs; also called habitat or ecological niche models) which estimate the potential biogeographic distribution of a species based on the environmental conditions it is observed in (Peterson & Soberón, 2012). Over the last two decades, SDMs have gained tremendous interest in the context of marine spatial planning (Melo-Merino et al., 2020). Initially developed for species occurrence data (Phillips et al., 2006; Thuiller et al., 2009), SDMs are now also applied (i.e., although marginally used) to continuous biomass or abundance data (Clerc et al., 2024; Knecht et al., 2023; Waldock et al., 2022), and proportions, such as those retrieved from metagenomic reads (Schickele et al., 2023). They benefit from numerous computational methods and guidelines transferred from other fields of research, that enhanced data integration, and led to a more robust assessment of their predictive capabilities, explicit uncertainties and quality control guidelines (Descombes et al., 2022; Leroy et al., 2018; Roberts et al., 2017; Waldock et al., 2022). However, these novel developments also led to a multitude of frameworks, each coming with their specific data requirements, formatting, quality checks, output format and limitations.

Therefore, there is a need for a common comprehensive SDM framework to produce high-quality, near real-time, standardized, and comprehensive EOVs and EBVs, as acknowledged at the international level (IPBES, 2016; IUCN Standards and Petitions Subcommittee, 2017). However, associating each output with explicit, multi-metric and community-approved quality control and metadata is still challenging. For instance, multiple taxonomic backbones are used or even developed, the associated metadata is highly heterogeneous and inherent data limitations remain, such as the management of heterogeneous sampling effort, biases in abundance measurements and the multivariate nature of omics data. Therefore, standard datasets, metadata and operative frameworks that emerged, or are used in the terrestrial realm (e.g., Feng et al., 2019; Fick & Hijmans, 2017; Hao et al., 2019; Wieczorek et al., 2012), are easily transferable to the specificities of the marine environment (Melo-Merino et al., 2020).

For the first time, we combine existing methods for presence only (Benedetti et al., 2021), continuous (Knecht et al., 2023), and proportion data (Schickele et al., 2023) in a common SDM framework for the marine environment, our new Comprehensive Ensemble Pipeline for Habitat modelling Across large-scale Ocean Pelagic Observation Datasets (CEPHALOPOD). Its novelty lies in the production of inter- comparable biogeographical estimates, EOVs, and EBVs, across multiple data types. To do so, CEPHALOPOD extrapolates information embedded in scarce and biased marine datasets and it is associated with comprehensive, harmonized, and explicit quality checks. Moreover, it directly accesses – but is not limited to – a large and updated collection of fish and plankton datasets and a new collection of 47 standardized environmental predictors. Here, we first estimate the performances of CEPHALOPOD using a virtual ecologist approach (Zurell et al., 2010). Then, we demonstrate its potential for extracting and mapping taxonomic diversity across data types through a case study applied to coccolithophore (i.e., unicellular calcifying phytoplankton species; Not et al., 2012) diversity patterns (O’Brien et al., 2016).

## MATERIAL AND METHODS

### 2.1. Overview of the workflow

The pipeline first accesses *in-situ* observation within – but not limited to – the Ocean Biodiversity Information System (OBIS; http://obis.org), Global Biodiversity Information Facility (GBIF; http://gbif.org), AtlantECO (Vogt et al., 2023), and Marine Atlas of Tara Ocean Unigenes (MATOU; Carradec et al., 2018) datasets (**Fig. 1**, steps 1-3). After selecting the taxa of interest, *in-situ* observations are associated environmental variables known to influence plankton biogeography (**Fig. 1**, step 4; **Table S1**). Both biological (target) and environmental (features) data undergo a series of common pre-processing and quality checks before model fitting. To avoid overparameterization, CEPHALOPOD constructs a set of non-collinear and informative environmental features (**Fig. 1**, step 6), associated to a quality flag (see *section 2.4*.; **Fig. 1**, step 6). Finally, the hyperparameters used for model tuning are defined and the training and testing datasets are built (**Fig. 1**, steps 7 and 8). At this stage, CEPHALOPOD integrated, processed, and quality-checked all input data and parameters necessary to calibrate an SDM algorithm (Feng et al., 2019).

**Figure 1:**
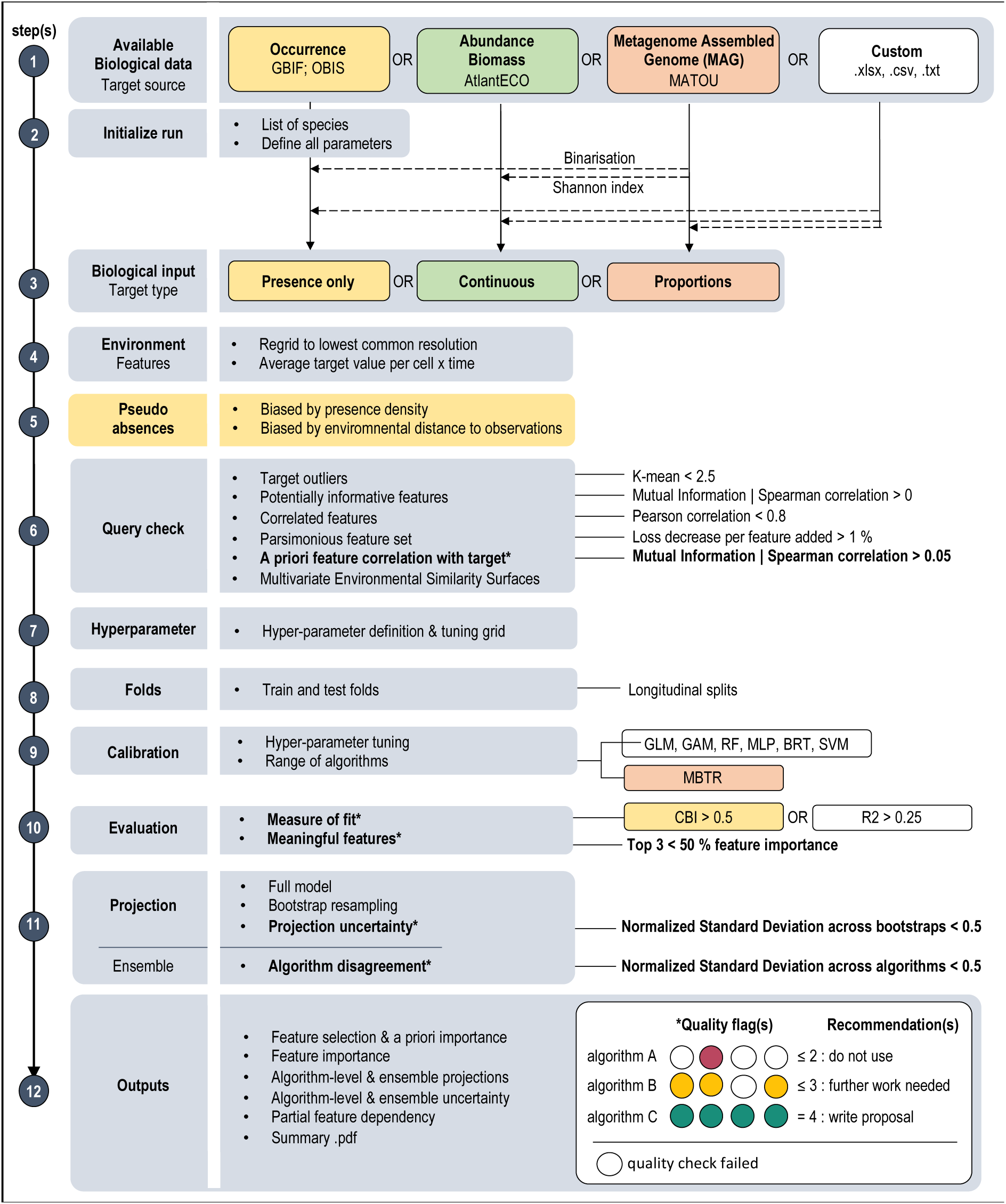
Synthetic diagram of the CEPHALOPOD indicating the essential list of modelling choices, quality checks and flags used in the workflow. The main steps are indicated on the left-hand side. The associated quality checks or modelling choices are displayed in the middle panels. The corresponding thresholds used for quality flags are indicated on the right-hand side. Those used in the final traffic-light quality flag are indicated in bold. Gray background corresponds to common steps or wrappers around target-type specific functions. Steps or functions are coloured in yellow, green, or orange if data-type specific.

CEPHALOPOD relies on an ensemble modeling approach, meaning it prioritizes spatial distribution estimates constructed from several algorithms. First, it selects the best hyperparameters for each algorithm using a spatial block cross-validation procedure (**Fig. 1**, step 9)(Roberts et al., 2017). Each algorithm output is then evaluated on the same cross-validation splits to estimate its performance in reproducing observations. Furthermore, we compute the importance of each feature in explaining these observations (**Fig. 1**, step 10). Both the model predictive performance and the interpretability of the feature importance are associated to a quality flag (**Fig. 1**, step 10). For all combinations of algorithms and taxa passing these quality controls, CEPHALOPOD performs spatial projections following a bootstrap procedure, to account for projection uncertainty (**Fig. 1**, step 11). We perform a final quality control on the projected uncertainty associated with each algorithm and across algorithms composing an ensemble projection (i.e., ensemble agreement; **Fig. 1**, step 11). Finally, CEPHALOPOD provides graphical outputs for each critical step and structured objects containing all inputs and outputs, directly usable for further computing. The quality checks associated with each set of observations are represented in the form of a traffic light and associated recommendations (**Fig. 1**, step 12).

### 2.2. Data preparation

CEPHALOPOD extracts relationships between presence only, continuous, or proportional biological target variables and environmental conditions. To this end, we need to (i) access both *in-situ* biological observations and (ii) environmental climatologies, (iii) retrieve and harmonize the corresponding relevant metadata (i.e., including taxonomy) and (iv) ensure parsimonious inputs (i.e., discard biological outliers or non-informative environmental climatologies).

#### 2.2.1. Biological target datasets

CEPHALOPOD accepts *in-situ* observations from different sources (**Fig. 1**, step 1 to 3), including a dynamic access to (i) occurrence data from OBIS and GBIF, (ii) abundances or biomasses retrieved from the AtlantECO dataset (Vogt et al., 2023) and (iii) metagenomic reads from MATOU (Carradec et al., 2018). To widen the range of possible applications, CEPHALOPOD accepts any target variables (e.g., other taxonomic records, biodiversity indices) that follow the Darwin Core format (Wieczorek et al., 2012), relevant for marine biogeographical data. CEPHALOPOD then filters spurious coordinates (e.g., 0°N – 0°E, land observations) and discards preserved specimen in museum collections, or fossil records. Finally, CEPHALOPOD matches the scientific names and associated identifiers against the latest taxonomy available in the World Register of Marine Species (WoRMS Editorial Board, 2024). When relevant, it updates the scientific names and identifiers to the accepted one and merges observations with synonym taxonomic annotations. Expert-based taxonomic aggregation is also possible by importing the target directly in CEPHALOPOD (**Fig. 1**, step 1, custom target).

#### 2.2.2. Matching environmental features with the target

CEPHALOPOD accesses a collection of observation-based monthly climatologies of potential environmental features (**Table S1**; **Fig. 1**, step 4) that were interpolated on a common grid (i.e., 1 x 1 degree resolution, global scale, gap-filled; Boyer et al., 2018). We consider monthly surface (0-50 m) or mesopelagic (200-300 m) climatologies encompassing the 1900-2020 period (33 out of 50 climatologies are post 1982). These climatologies characterize the physical (e.g., sea surface temperature), chemical (e.g., pH) and biological (e.g., Chlorophyll-a concentration) conditions of water masses as well as their circulation and turbulence patterns (e.g., Eddy Kinetic Energy; EKE). All these features have been shown to have an influence on the biology and spatial distribution of marine pelagic organisms (Benedetti et al., 2021; Dahlke et al., 2020; e.g., Melo-Merino et al., 2020). Finally, the target is gridded on the spatio-temporal grid of the features and the values are averaged to decrease sampling bias.

#### 2.2.3. Pseudo absences selection for presence only data

When using presence only data, we need to compensate the lack of true absences by generating pseudo-absences to train regressive SDMs. The pseudo-absence choice is a key uncertainty in SDM modelling, and should be subject to the same sampling bias as the presence data (Descombes et al., 2022; Righetti et al., 2023). Therefore, CEPHALOPOD samples pseudo-absences following by default the nearby density of presences (i.e., Gaussian focal, radius σ = 20; **Fig. 1**, step 5) and an environmental envelope (i.e., environmental conditions outside the range of presences; Getz & Wilmers, 2004; Schickele et al., 2020). Other pseudo-absence selection methods – and combinations thereof – can be selected, including a minimum geographical distance to presences, or a user-defined sample probability background (i.e., for target group approaches; Righetti et al., 2023). For each biological target, CEPHALOPOD samples pseudo-absences in equal number and weight to the presences, to minimize the prediction variance (i.e., D-design theory; Montgomery, 2017).

#### 2.2.4. Final pre-processing and quality check

For each target variable, CEPHALOPOD performs a series of quality checks and pre-processing steps prior to training the SDM algorithms (**Fig. 1**, step 6). First, outliers in the target are discarded according to the Z-score criterion, defined as a 2.5 standard deviation from the mean target value across observations (e.g., Knecht et al., 2023). Second, environmental features that are unlikely to explain the target distribution are discarded: CEPHALOPOD discard features that present a higher spearman correlation or mutual information with a null target (i.e., random values sampled within the range of the biological target) than the biological target. To avoid variance inflation, we assess multicollinearity by performing a pairwise correlation of the features at the target location (F. Dormann et al., 2007). Among each group of collinear features (Pearson r > 0.8), we only considered the one sharing the highest correlation or mutual information with the target. Yet, this feature selection may still contain duplicated information due to the multivariate aspect of the problem. Therefore, CEPHALOPOD then performs a Recursive Feature Elimination procedure (RFE; Darst et al., 2018; R library “caret” version 6.0-93; non-applicable to proportion data due to their multivariate nature). This sequentially (i) fits a Random Forest algorithm using all features and a 5-fold cross-validation procedure, (ii) computes a loss between observed and predicted values (R^2^), (iii) ranks the features by their predictive importance (%) and (iv) discards the least important one. As a result, CEPHALOPOD ranks the features by their *a-priori* importance and selects the number after which adding a supplementary feature does not decrease the loss (i.e., calculated as a moving average of size 5). Finally, to identify the regions where SDM extrapolate their predictions into non-analogue conditions, CEPHALOPOD performs a Multivariate Environmental Similarity Surfaces (MESS; Elith et al., 2010).

### 2.3. Predicting the distribution of the biological target

#### 2.3.1. Training design and algorithms

To account for uncertainties related to algorithm choice, we use an ensemble approach as recommended by Hao et al. (2019). For presence only and continuous data types, CEPHALOPOD fits six algorithms across a wide spectrum of statistical and machine learning tools, ranging from simple to complex models: Generalized Linear Model (GLM), Generalized Additive Model (GAM), Multilayer Perceptron (MLP), Random Forest (RF), Boosted Regression Tree (BRT) and Support Vector Machine (SVM). For proportional data, CEPHALOPOD fits a Multivariate Boosted Tree Regressor (MBTR), a recently developed algorithm specific to multivariate data (Schickele et al., 2023). To minimize both overfitting and the effect of spatial autocorrelation which may lead to overestimated performance estimates (Roberts et al., 2017), CEPHALOPOD splits the target and feature datasets between training and testing sets using a *n*-fold cross-validation procedure and spatial blocking according to the longitudinal coordinates of the samples (**Fig. 1**, step 8). Each algorithm is trained *n*-times on different *n*-1 folds, while the remaining fold is kept for testing only. Therefore, an algorithm overfitting the training sets results in lower predictive performance on spatially independent test sets. First, CEPHALOPOD estimates the best hyperparameters across the *n*-cross validation folds for each algorithm and target (**Fig. 1**, step 9). Using the best hyperparameters, CEPHALOPOD then estimates the predictive performance of each algorithm by predicting the same *n*-test split (see *section 2.4*., **Fig. 1**, step 10).

#### 2.3.2. Spatial projection and uncertainty

To estimate the projection uncertainty associated with each species and algorithm, CEPHALOPOD performs a 10-time bootstrap resampling procedure (**Fig. 1**, step 11). For each bootstrap round, it trains a full model by using the best hyperparameters estimated in **Fig. 1**, step 9 (*section 2.3.1*.) on a resampling of the original dataset (i.e., length equal to the original dataset, with replacement). Then, CEPHALOPOD projects the corresponding monthly habitat suitability on the global scale, using the feature values associated to each grid cell. All algorithms that successfully pass the quality checks are aggregated in an ensemble projection (see *section 2.4*.; **Fig. 1**, step 11).

#### 2.3.3. Derived outputs

CEPHALOPOD offers the possibility to compute partial dependence plots for each feature included in the SDMs (**Fig. 1**, step 12), i.e., a mapping of the marginal response of the biological target to each environmental feature in environmental feature space. Partial dependence plots are crucial outputs to interpret the ecological response of each species to the environmental gradients and discuss the underlying biological mechanisms, as well as to diagnose over-fitting. Finally, CEPHALOPOD can compute a range of diversity indices across multiple taxa, such as the Shannon, Richness and Evenness indices to highlight biodiversity hotspots across the global ocean (Magurran, 2011).

### 2.4. Quality assessment and flags

We designed CEPHALOPOD around a multi-metric SDM evaluation that integrates four major and standardized quality flags. First, the *a-priori* importance of the considered environmental features in explaining the biological target is tested by their average shared spearman correlation and mutual information (Kinney & Atwal, 2014), which must be 0.05 higher than the one with a null target (**Fig. 1**, step 6) (i.e., adapted from Knecht et al., 2023). Second, CEPHALOPOD estimates the predictive performance of each algorithm by quantifying the quality of the corresponding evaluation split prediction (**Fig. 1**, step 10). To do so, CEPHALOPOD computes (i) a Continuous Boyce Index (CBI; Hirzel et al., 2006), (ii) a univariate r-squared (R^2^; Pearson), or (iii) a multivariate r-squared (Pearson) for presence only, continuous and proportional data, respectively. All metrics range from -1 to 1 and correspond to the closest equivalent, adapted to each data type. The pipeline only considers algorithms presenting a predictive performance above 0.5 for presence only data, or 0.25 for continuous and proportional data (**Fig. 1**, step 10). Third, CEPHALOPOD extracts the importance of each feature in explaining the observed patterns in the target (i.e., see function specification in the “tidymodels” R library, version 1.1.1.). To better interpret underlying ecological processes, informative features should explain a substantial part of the observed variability. Thus, CEPHALOPOD retains only algorithms with a cumulative variable importance above 50% for the top three features (**Fig. 1**, step 10). Algorithms successfully passing the above-mentioned quality checks are retained to perform habitat suitability projections at the global scale. Finally, CEPHALOPOD computes the scale independent Normalized Standard Deviation (NSD), defined as the standard deviation normalized by the maximum average across bootstrap runs, for each algorithm, month, and grid cell. To quantify the global projection uncertainty for each algorithm, the pipeline averages the NSD across the geographical projection domain and associates a quality check: CEPHALOPOD retains only algorithm with an NSD averaged across grid cells, under 0.5 (**Fig. 1**, step 11). The pipeline also computes the NSD between each algorithm considered in the ensemble projection as a proxy for disagreement among algorithms. CEPHALOPOD retains only ensembles with an NSD across algorithms under 0.5 (**Fig. 1**, step 11). The above-mentioned sequential quality checks are displayed in a comprehensive traffic-light system with associated recommendations. For each algorithm or ensemble, CEPHALOPOD displays a red, yellow, or green light highlighting successful quality checks, with the colour corresponding to a success for at least 2, 3 or 4 quality checks, respectively (**Fig. 1**, step 12, **Fig. S6**).

### 2.5. Applications

#### 2.5.1 Testing CEPHALOPOD with virtual species

We evaluated the skill of CEPHALOPOD in reproducing a set of simulated species distributions across presence only, continuous and proportional data through a “virtual ecologist approach” (Zurell et al., 2010) using the “virtualspecies” R library (version 1.6; Leroy et al., 2016). First, we perform a Principal Components Analysis (PCA) from all the environmental features available. Then, we create a virtual niche by constructing a normal distribution along the first and second PCA axis (i.e., with µ and σ sampled randomly within the corresponding PCA axis space). We simulated a set of 20 virtual niches, resulting in spatial distributions encompassing prevalences (i.e., the proportion of geographical cells where the species is considered present) ranging from 0 to 1. This way, we mimic the distribution of both ubiquitous and rare species. We then sampled these 20 simulated distributions, assumed as ground truth. To generate discrete presence records from the continuous habitat suitability maps (where values are in [0,1]), we performed a binomial trial centered on 0.5 in every virtual sampled cell; if the value of the binomial trial was lower than the habitat suitability, then the species was considered present in this cell. We generated samples of three different sampling efforts (N = 50, 200, 1000). For each sampling effort, we generated samples of three geographical bias levels, either (i) homogenously across the global ocean, (ii) with a coastal or (iii) north Atlantic bias to mimic commonly found spatial biases (**Fig. 2**). For continuous and proportional data, we also added different levels of background noise to the sampled values (i.e., log-normal bias with standard deviation of 1 and 2; **Fig. 2**). Doing so, mimic the variability induced by local processes and measurement error. This virtual species design results in 180 and 540 datasets for the (i) presence only and (ii) continuous or proportion target types, respectively.

**Figure 2:**
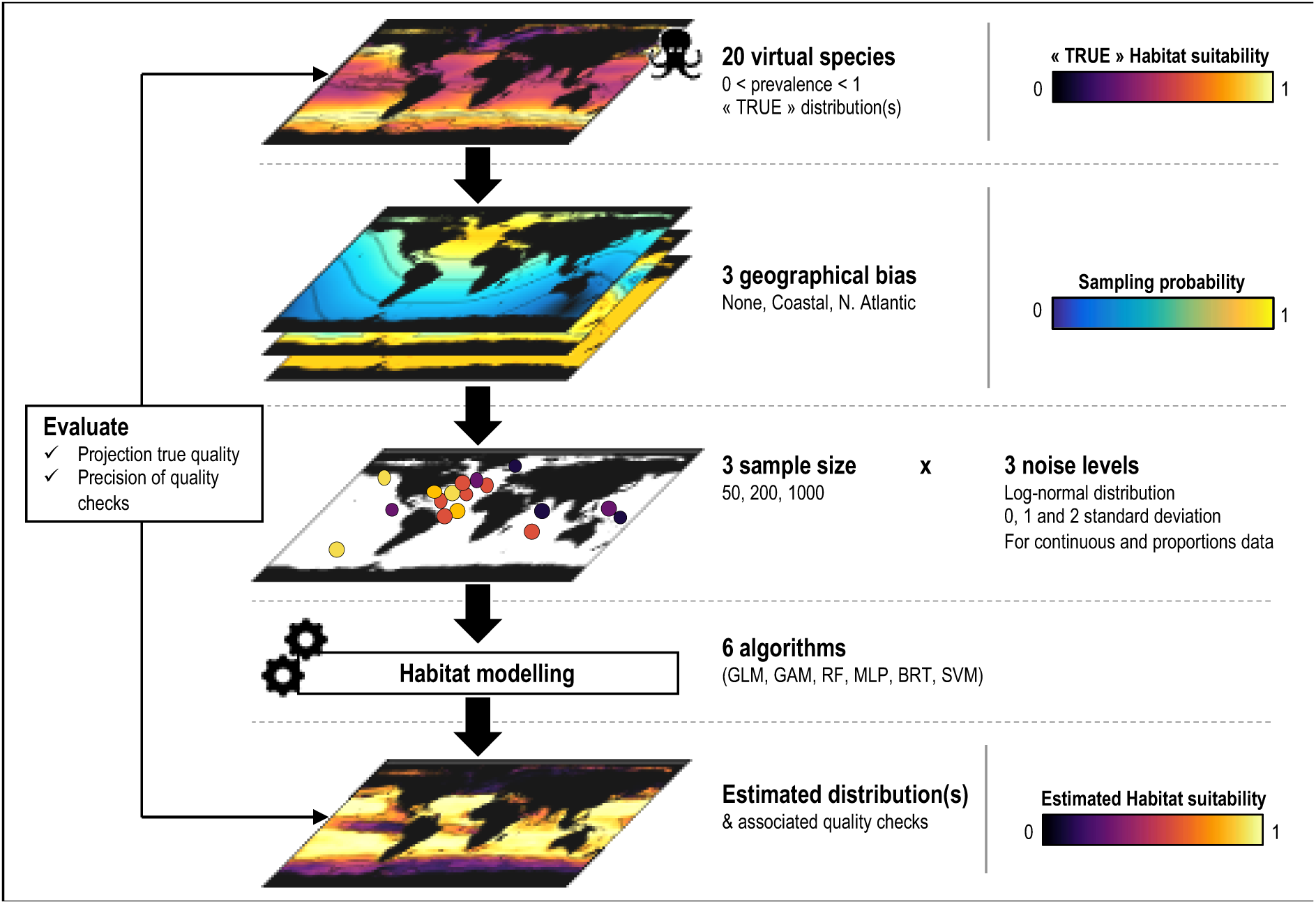
Synthetic diagram of the framework testing on virtual species. The virtual distributions, geographical biases, sample sizes and noise levels are indicated in lines.

#### 2.5.2. Application to coccolithophores diversity from real observations

Furthermore, we present a case study based on in-situ observations by investigating coccolithophores (i.e., *Prymnesiophyceae*) diversity patterns across data types in the global epipelagic ocean. First, we retrieved *in-situ* observations of coccolithophores abundance from AtlantECO-base (Vogt et al., 2023). Following the methods described in O’Brien et al. (2016), we integrated coccolithophores observations across geographical cells, month, and surface ocean, from which we computed a richness estimate by using a 100-times rarefaction procedure (i.e., each sample is resampled 100 times with replacement to alleviate the effect of different sample size). This procedure is known as an “assemble first – predict later” diversity estimation (i.e., diversity is computed from the observations and modelled as a target in the SDMs). We estimated the global scale coccolithophores richness using a similar selection of environmental features as in O’Brien et al. (2016).

Second, we estimate coccolithophores Shannon diversity from Metagenome Assembled Genomes (MAGs) based on relative genome reads from the MATOU catalogue among the 0.5 – 8 µm size filter, corresponding to the *Prymnesiophyceae* size range (Not et al., 2012). Furthermore, MAGs are considered as species (*sensu* metagenomics) and their definition is freed from the taxonomic annotation and harmonization issues. We standardized each read by the gene length and sequencing depth, ensuring comparable community samples across Tara Ocean stations (Schickele et al., 2023). We computed a coccolithophores Shannon index estimate from the resulting proportions of metagenomic reads (“assemble first – predict later”). Finally, we used CEPHALOPOD to compute spatial projections of the relative metagenomic reads, treated as proportional biological targets. Doing so, we also computed a coccolithophores Shannon index from the projections (i.e., “predict first – assemble later” approach; the diversity is computed on each cell of the global ocean). We also estimated a Shannon index based on abundances using the method described above. We estimated all global scale coccolithophores Shannon indices using the full set of available environmental epipelagic features (**Table S1**).

## RESULTS

### 3.1. The predictive power of CEPHALOPOD

To test whether CEPHALOPOD produces reliable distribution estimates across data types and sample quality, we first evaluated its ability to reproduce virtual species distributions, as discussed in *section 2.5.1*., based on a range of derived samples across combinations of prevalence, size, geographical bias, and background noise (**Fig. 3A-C**). The resulting projections, successfully passing our four quality checks, present a median Pearson correlation with the true distribution of 0.79, 0.82 and 0.58 for presence only, continuous and proportions data, respectively (**Fig. 3A-C**). The associated standard deviation across all grid cells expressed as median [inter-quartile range] relative to their corresponding true distribution, is of 1.23 [1.10 -1.49], 1.06 [0.87-1.23] and 1.0 [0.75-1.28] (**Fig. 3A-C**). This means that CEPHALOPOD produces distribution estimates whose spatial pattern and variability are consistent with the corresponding true distribution. This consistency is valid all data types and sampling strategies. To further investigate the skill of CEPHALOPOD, we performed an analysis of variance (ANOVA) to identify which factors affect the model performance in predicting the true distribution. We show that prevalence, geographical bias, background noise, sample size, and algorithm choice all show a significant effect on true predictive performance across most data types (**Fig. S2** and **S3**). However, the relative contribution of each factor highlights only a major contribution of prevalence (53%) when using presence only data and geographical bias (45 and 46%) when using continuous or proportional data (**Fig. S2**). This means that CEPHALOPOD may produce lower quality (i.e., although still consistent) distribution estimates when using presence only data for ubiquitous species (i.e., those with prevalences > 0.8; Pearson correlation of 0.56; **Fig. 3A**) or when using continuous or proportional data for regionally biased samples (i.e., north Atlantic bias; Pearson correlation of 0.73 and 0.26, respectively; **Fig. 3B**-**C**).

**Figure 3:**
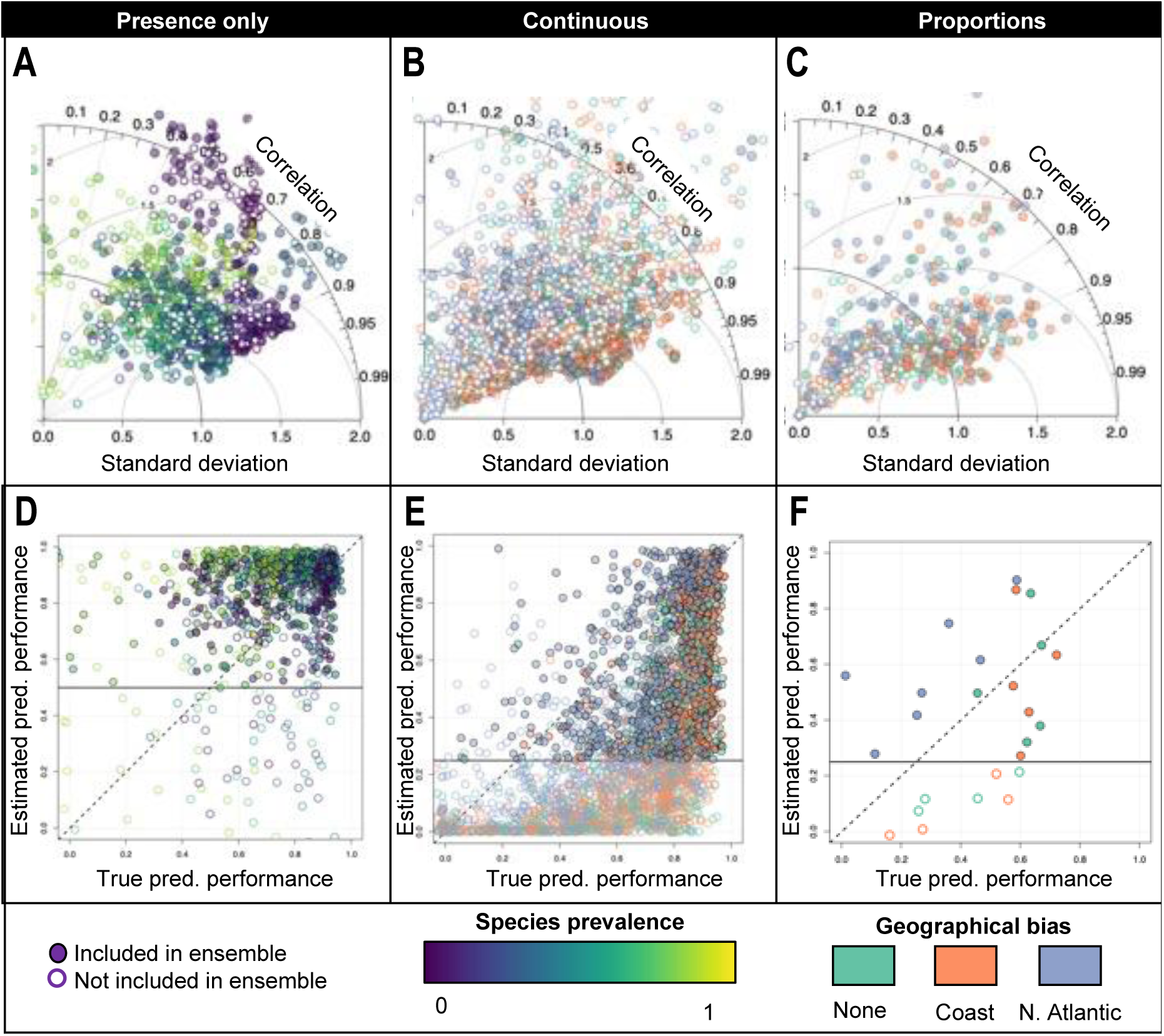
Habitat modelling performance on 20 virtual species for presence only (**A**, **D**), continuous (**B**, **E**) and proportion data (**C**, **F**). Each circle corresponds to a combination of prevalence, sample size, background noise, geographical bias, and algorithm. For each data type, the circles are coloured by the factor explaining the most variability. The top panels (**A**-**C**) compare the estimated and true distribution. The bottom panels (**D**-**F**) compare the estimated predictive performance (i.e., observed vs predicted evaluation split) to the true performance (i.e., estimated vs true distribution). The 1:1 dashed line indicates a perfect performance estimation. The horizontal black line indicates the threshold under which an algorithm is discarded. Note that the estimated predictive performance is computed simultaneously across all targets due to the multivariate nature of proportion data (**F**).

To evaluate whether CEPHALOPOD accurately reports the true distribution estimate quality, we compare the true predictive performance of each algorithm against the one estimated by the corresponding quality checks (i.e., CBI or R^2^; see *section 2.4*. and **Fig. 1**, step 10). These quality checks present median values across each algorithm and sampling strategy of 0.91, 0.54 and 0.51 for presence only, continuous, and proportional data, respectively (**Fig. 3D-F**). The associated deviation between the true predictive performance and the one estimated by CEPHALOPOD present an RMSE of 0.27, 0.31, and 0.26 (**Fig. 3D-F**). This means that the predictive performance estimated by CEPHALOPOD is consistent or under-estimated (i.e., for continuous data; **Fig. 3E**), and shows a limited deviation, compared to the true predictive performance. To further investigate the robustness of our quality checks, we performed an ANOVA to identify which sampling strategy affects the false positive rate in the estimated predictive performance. We show that prevalence, geographical bias, background noise, sample size, and algorithm choice all show a significant effect across most data types (**Fig. S4** and **S5**). However, the relative contribution of each factor highlights only a major contribution of prevalence (73%) when using presence only data and geographical bias (34 and 43%) when using continuous or proportional data (**Fig. S4**). This means that potential false positive quality flags may relate to the use of presence only data for ubiquitous species (**Fig. 3D**) and continuous or proportional data for regionally biased samples (i.e., north Atlantic bias; **Fig. 3E**-**F**).

Overall, our virtual ecologist approach shows that each algorithm considered in CEPHALOPOD performs well at reproducing distributions based on samples of prevalence, size, geographical bias, and background noise from biased samples (median Pearson correlation of 0.72; **Fig. 3**). This performance is accurately estimated by CEPHALOPOD, with a limited risk of false positive quality flags on the distribution estimates (**Fig. 3**).

### 3.2. Overview over standard outputs of CEPHALOPOD

Here, we first present a selection of outputs as a key example of our new modelling pipeline, corresponding to the ensemble estimation of coccolithophores richness (see *section 2.5.2*.) The first key output of CEPHALOPOD is the traffic-light evaluation system (**Fig. 4A** and **S6**), performed for each biological target variable and algorithm. Here, the pipeline returns a green light for the ensemble estimation as all quality checks are successful (see *section 2.4*. and **Fig. S6**). The environmental features selection and their interpretability are satisfying, the algorithms considered showed good performance in reproducing the observed patterns, with moderate projection uncertainty, leading to positive recommendations (**Fig. 4A**). Temperature (27 %) emerges as the main feature explaining the spatial distribution of coccolithophores diversity, followed by phosphate concentration (26 %) and silicate concentration (20 %; **Fig. 4B**). The resulting ensemble projection (**Fig. 4C**) matches the estimates of O’Brien et al. (2016): we find high coccolithophores richness (> 10) predicted in the equatorial Pacific and Indian oceans, excluding the east pacific equatorial upwelling, and lower richness (< 3) at higher latitudes (> 50°). The hotspots of annual coccolithophores richness (Q75; **Fig. 4D**) overlap with observations in the equatorial and tropical Indian ocean, whereas low richness estimates overlap with the corresponding observations in higher latitudes. Here, the ensemble agreement present intermediate values (NSD of 0.21), with higher values in between 50°N and 50°S. Finally, the low MESS values indicate a good environmental coverage of the observations (**Fig. 4E**).

**Figure 4:**
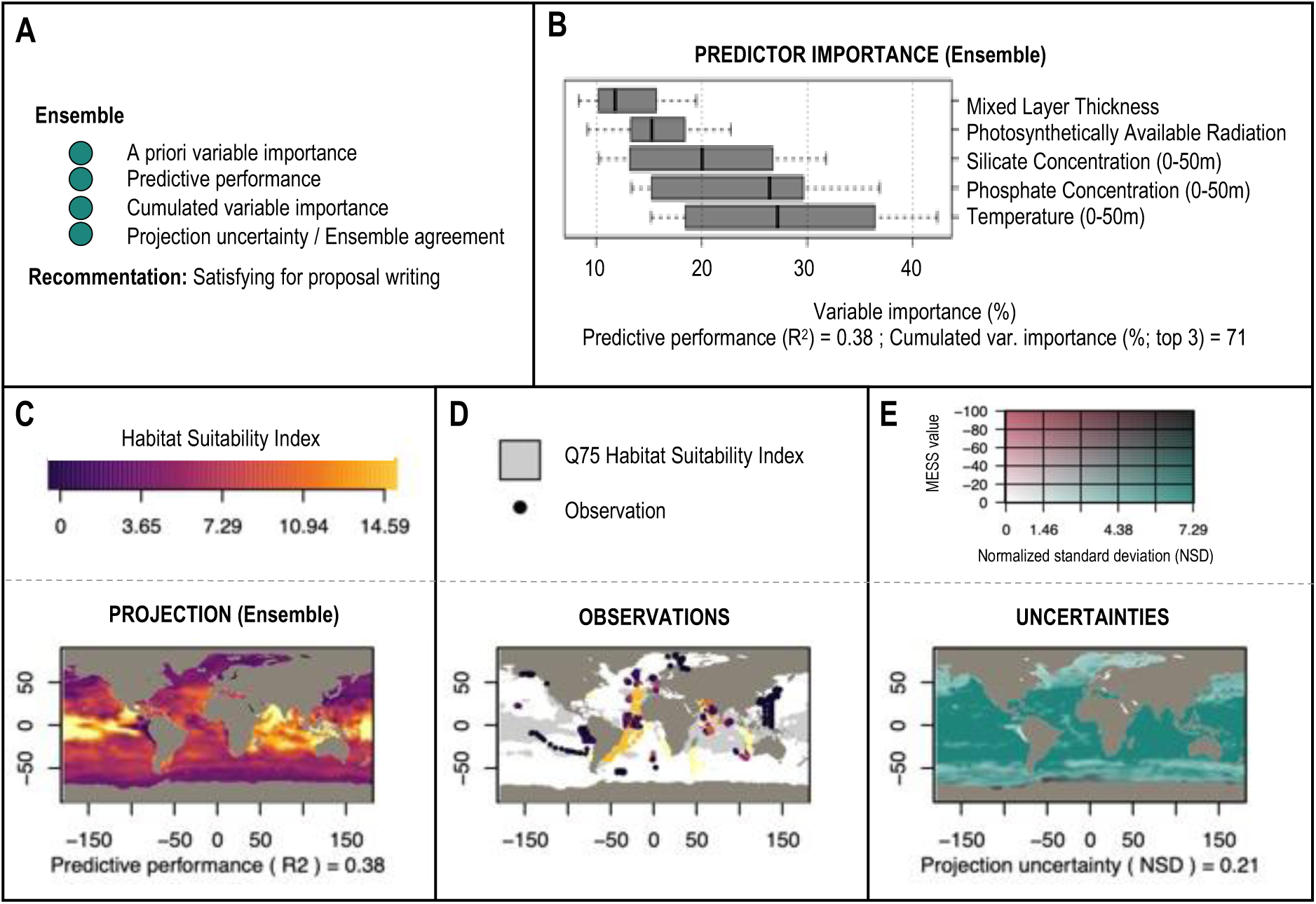
Overview of a default output from CEPHALOPOD including the (**A**) traffic light quality check, (**B**) environmental predictor importance, (**C**) spatial distribution projection, (**D**) observations and (**E**) associated uncertainties. In (**D**), Q75 correspond to the 75^th^ percentile of the projected values. In (**E**), MESS corresponds to the multivariate environmental surface similarity, an estimation of extrapolation outside the environmental conditions of the observations. Note that labels are clarified in relative to the programmatic output.

### 3.3. Exploring global coccolithophores diversity across data types

We now show the mean annual patterns of global Coccolithophores Shannon diversity across data types and assembling methods (i.e., assemble first – predict later vs predict first – assemble later), estimated by CEPHALOPOD (**Fig. 5**).

**Figure 5:**
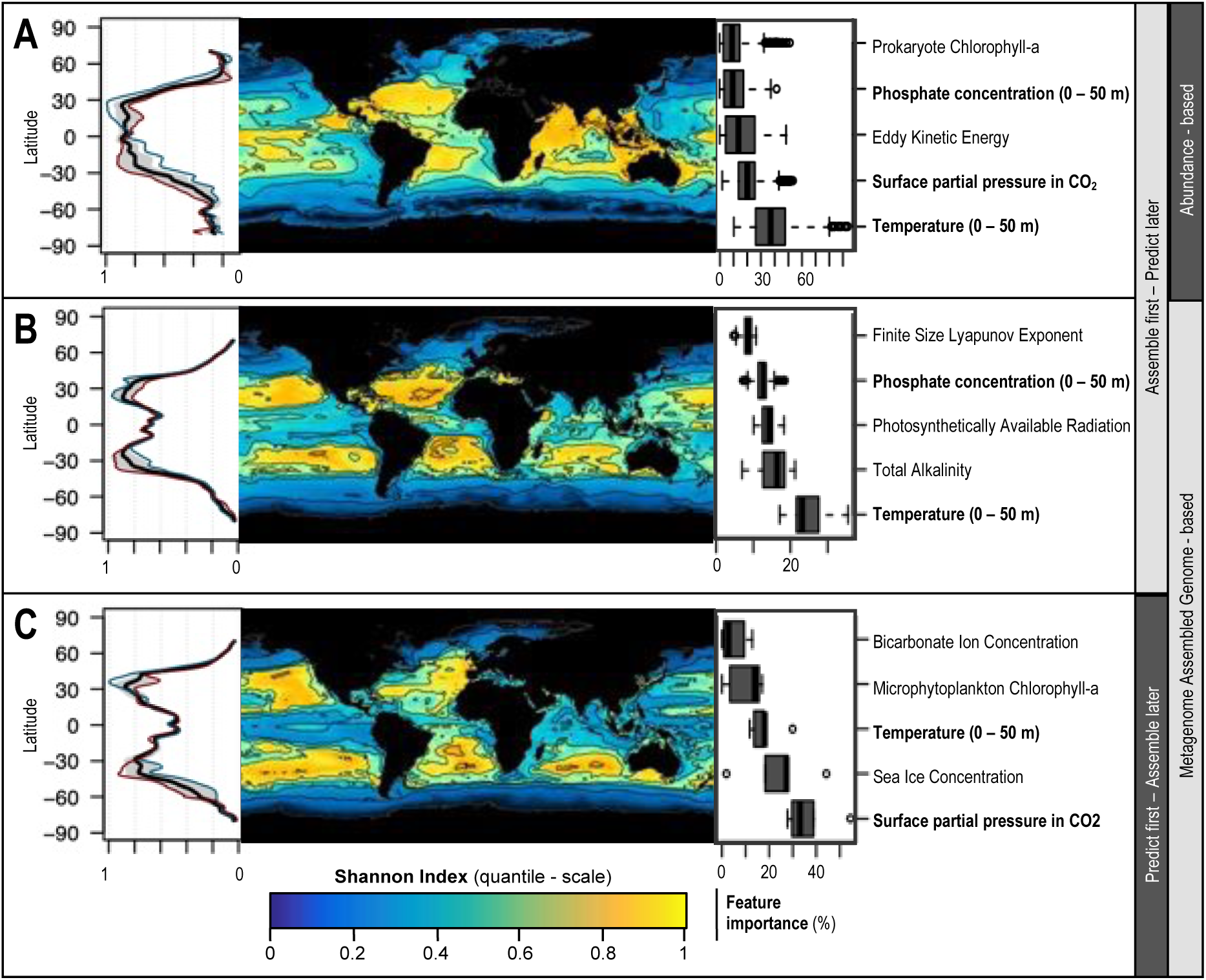
Coccolithophore Shannon index estimates across data types and methods, including assemble first – predict later from (**A**) abundance (i.e., continuous), and (**B**) metagenome assembled genomes (MAGs; continuous), and predict first – assemble later from MAGs (i.e., proportions) The Shannon index projections are represented by the quantile distribution, averaged across all months. The latitudinal profiles show the corresponding average (black line), January (blue line), July (red line) and standard deviation (grey shading) across months. Important environmental features are highlighted in bold when identical across data types and methods.

Based on abundance data, the mean annual coccolithophores Shannon index shows a strong latitudinal gradient characterized by maxima in the tropical ocean and minima in the polar oceans (**Fig. 5A**). The tropical Atlantic and Indian oceans present values above the 0.8 quantile, which identifies them as hotspots of Coccolithophores diversity. Again, our results concur with the richness estimates from O’Brien et al. (2016) (**Fig. 3C** and **5A**), which suggests that CEPHALOPOD accurately reproduces ecological conclusions from previous studies. Moreover, the January and July profiles highlight an important seasonal variation in latitudes corresponding to tropical gyres (30°N and 30°S) with an estimated quantile standard deviation of 0.4 (**Fig. 5A**). Temperature (38 %) is the main environmental feature explaining the coccolithophore Shannon diversity from abundance, as in O’Brien et al. (2016).

Based on the metagenomic reads of MAGs, mean annual coccolithophores Shannon index also shows a strong latitudinal pattern with diversity hotspots in oligotrophic gyres when using both the assemble first (**Fig. 5B**) or the predict first (**Fig. 5C**) strategy (i.e., Spearman correlation coefficient of 0.85). The associated latitudinal patterns are bimodal and centred on the tropical gyres (30°N and 30°S). They also present a low seasonality as shown by the low monthly standard deviation and similarity between the January and July profiles (**Fig. 5B** and **C**, left panels). Finally, temperature and partial pressure in CO_2_ are shared important (> 20%; phosphate concentration in a lesser extent) features explaining the Shannon index patterns across both data types and assembling method (**Fig. 5**).

All Shannon index estimates exhibit successful quality checks (see *section 2.4*.) with (i) a predictive performance of 0.54, 0.67 and 0.58, (ii) a cumulated variable importance of 54, 76 and 79 %, and (iii) NSDs of 0.17, 0.03 and 0.41 for assemble-first from abundances, assemble-first from MAGs and predict-first from MAGs, respectively. This means that CEPHALOPOD not only successfully reproduces previous ecological findings but also produces robust distribution estimates associated with consistent environmental drivers from scarce observations and new types of data not previously modelled.

## DISCUSSION

### 4.1. Advantage of CEPHALOPOD

The most notable strengths of CEPHALOPOD are (i) its capacity to harmonize observed distributions, (ii) and its standardized quality flags across multiple data types. It also offers the possibility to perform inter-comparable projections of multiple biological target species, across several data types. Moreover, the traffic light recommendations comprehensively report on the reliability of model outputs and thus matches the requirements for minimum SDM standards (Flombaum & Martiny, 2021). We showed how CEPHALOPOD can successfully reproduces species diversity estimates from previous studies (O’Brien et al., 2016). Therefore, it enables a quantitative inter-comparison between original research findings and previous works. Furthermore, CEPHALOPOD is largely automatized and parallelized in its data pre-processing and modelling steps, therefore greatly facilitating the estimation of species distribution and diversity across a large range of taxa. Other SDM frameworks propose similar quality checks (e.g., predictive performance, uncertainty, and extrapolation quantification), modelling options (e.g., ensemble modelling, hyper-parameter tuning, pseudo-absence selection) and outputs (e.g., projections, variable importance). However, they are often specifically designed for one type of data (e.g., occurrences; Phillips et al., 2006; Thuiller et al., 2009), therefore limiting the possibilities of integration and inter-comparison across studies or outputs from other data types such as omics or quantitative imaging. To the best of our knowledge, CEPHALOPOD is the first available SDM framework targeted at multiple data types that proposes comprehensive, reproducible, and inter-comparable species distribution and diversity estimates.

### 4.2. Caveats and limitations

Nevertheless, certain limitations intrinsic to habitat modelling persist in CEPHALOPOD: the lack of factors integrating dispersal limitations (e.g., in case of invasive species), explicit species interactions or other life history traits influencing species distribution (e.g., diel vertical migrations). While these limitations are inherent to habitat modelling, they mostly relate to local scale processes, which are outside the scope of our framework (Holt, 2020). The automatized modelling steps, while objectively chosen in a mathematically consistent way, may not always lead to distribution estimates or features of ecological significance. For instance, selecting salinity lead to basin-specific biases when regions of higher salinity such as the North Atlantic Ocean are disproportionally sampled (Benedetti et al., 2021; Boyer et al., 2018). This is why we left the possibility for expert-based knowledge to guide features and parameters selection in each modelling step, even if CEPHALOPOD is designed for a high degree of automatization. In addition, the quantitative observations retrieved from traditional biomass or omics-based data are not always comparable among sampling techniques, due to differing capturability or completeness of the sample across net types, mesh sizes, or sequencing depth (Lombard et al., 2019). Divergent taxonomic nomenclatures also reduce comparability between data sources, particularly for prokaryotes, not yet included in the WoRMS. However, global diversity occurrence-based assessments should be independent of the individual taxonomic annotations or quantitative biases in the *in-situ* observations. To face these inherent limitations, CEPHALOPOD has been constructed on best available practices and provides explicit insights into its performance, biases as well as guidance towards informed decision-making in diverse ecological contexts.

### 4.3. Case studies

Our case studies offer additional insights on how well CEPHALOPOD performs compared to alternative modelling pipelines in previous work, and its ability to detect ecological processes from an inter- comparison application.

First, the virtual case study highlights the overall good performance of CEPHALOPOD across multiple data types and sampling biases common to marine biogeographical data (Hughes et al., 2021). It also highlights the value of CEPHALOPOD in identifying cases of poor model performance, which limits the risk of misuse or misinterpretation of the estimated distributions. Indeed, the virtual species approach suggests a tendency to overestimate the distribution quality when using presence only to model ubiquitous species, or for geographically biased samples. The former is a known long-standing issue in habitat modelling, as model evaluation is particularly difficult when the environmental conditions of a species’ presence overlap with those of the background data (Tsoar et al., 2007). These species may correspond to ubiquitous protists, some of which are major contributors to global taxonomic richness patterns (Chapman et al., 2018). Geographically biased sampling effort is also known to bias distribution estimates, when the environmental conditions of observations are not representative of the fundamental niche of a taxa, commonly known as niche truncation (Chevalier et al., 2022). Nevertheless, CEPHALOPOD provides a robust quality assessment that enables inter-comparable distribution estimates from various data types, its main objective.

Second, the real case study shows that diversity estimates from abundance and MAGs present different patterns, despite similar environmental drivers. Indeed, different types of data shed light on different ecological processes that influence distribution estimates, and the associated diversity (Waldock et al., 2022). For instance, net-based plankton abundance measurements provide insights into population size or growth (Dallas & Hastings, 2018; Waldock et al., 2022), while, metagenomic reads relate to community composition and evolutionary processes (Delmont et al., 2022; Obiol et al., 2020). Despite these differences, temperature, and nutrients concentrations, emerged as major drivers of coccolithophores diversity out of a collection of 47 environmental features. Both are known to shape phytoplankton distribution (e.g., O’Brien et al., 2016; Righetti et al., 2019; Zhong et al., 2020), supporting the ability of CEPHALOPOD to select relevant features and response curves. Conversely, we found converging coccolithophores diversity pattern between different assembling methods, despite diverging environmental drivers. This suggests that coccolithophore diversity is independent of the assembling method and may therefore be an emergent property of the system. Therefore, CEPHALOPOD also encourages the exploration of ecological processes that arise from the interplay between different data types and assembling methods.

## CONCLUSION

CEPHALOPOD pushes beyond traditional habitat modelling tools by embracing the multidimensionality and the diversity of data types generated to study marine ecosystems, while making them inter-operable and inter-comparable. As an algorithm with a high degree of automatization potential, it is particularly suitable for the direct coupling with international data repositories (Martín Míguez et al., 2019; Mitchell et al., 2020) within the Digital Twin of the Ocean framework. The flexible and standardized formatting of *in-situ* observations datasets in CEPHALOPOD supports the development of FAIR datasets including biovolume, biomass or traits from quantitative imaging, or various taxonomic and functional resources derived from cutting-edge omics methods (Dugenne et al., 2023; Lombard et al., 2019; Tanhua et al., 2019). To this end, CEPHALOPOD has the potential to address evolving demands of multidimensional ecological research (Ashcroft et al., 2017; Mostert & O’Hara, 2023). By producing inter-comparable EOVs and EBVs, CEPHALOPOD also offers an operational tool of interest for a broad audience across both scientific, policy and conservation communities.

## STATEMENTS

## Acknowledgements

The authors wish to thank public taxpayers who fund their salaries. A.S., C.C., F.B. D.D.A. and U.H.E.’s salaries were financed by the BIOceans5D, Bluecloud2026 and AtlantECO European projects. The authors want to thank all people involved in the Tara Oceans and AtlantECO projects for making data publicly available. M.V would like to thank N. Knecht, L. Gregor for the conception of a first prototype during the first Bluecloud hackathon. M.V., F.B. and D.D.A. would like to acknowledge L. Maiorano and the AtlantECO WP2 team for data provision and model expertise. A.S., C.C., M.V. and U.H.E. would like to thank S. Pesant, R. Finn, L. Richardson, L. Meszaros, D. Obaton and the Bluecloud scientific and technical coordination team for fruitful discussion and advice on data structures and cloud-computing. A.S., C.C., and M.V. would like to acknowledge Cael B.B., G. Chust and the B5D modelling community for valuable input on modelling standards during a recent workshop. A.S., C.C., M.V. and U.H.E. would like to thank S. Lunven for extensive testing of the pipeline.

## Authors contribution

M.V. conceived and supervised the study and provided fundings. A.S., C.C. and F.B. processed the input and predictor data and provided expert input. A.S. wrote the first draft, conceived, and designed the modelling pipeline and performed the analysis. J.O.I. and D.D.A. provided expertise on SDM modelling across data types. All authors substantially contributed to the successive versions of the modelling pipeline and of this manuscript.

## Data availability

The primary biological data supporting the conclusions of this manuscript are available at https://zenodo.org/records/7944433 and https://data.d4science.net/qa7Z. The collection of environmental predictors is available at: https://data.d4science.net/m9WC. We intend to archive all other data associated with this manuscript in Zenodo where applicable.

## Code availability

The code supporting the conclusions of this manuscript are available in the following repository: https://github.com/alexschickele/CEPHALOPOD

