## Supplementary material for "Standardizing marine habitat modelling practices to enhance inter-comparability across biological observations": Fig. S

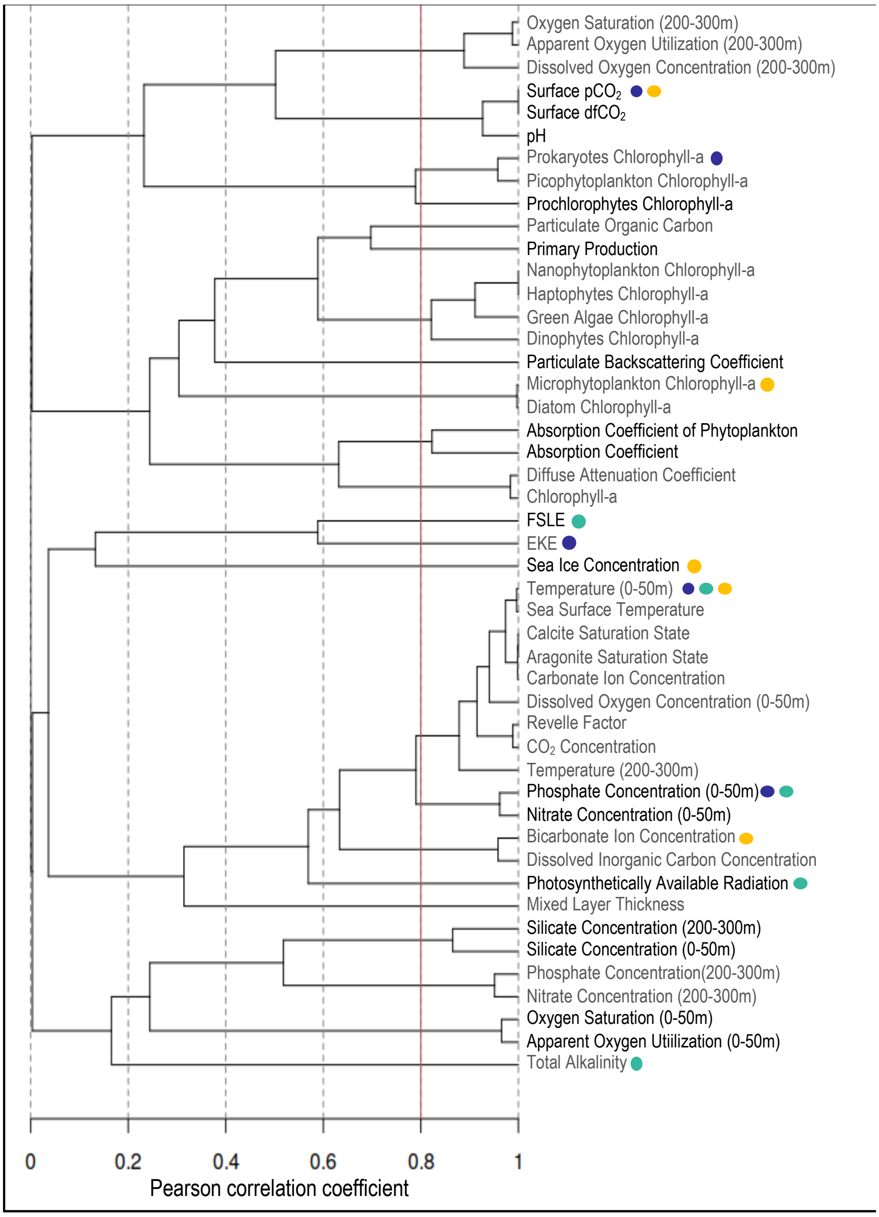


**Figure S1**: Pearson correlation coefficient and associated hierarchical clustering between environmental predictors (i.e., features) at the global scale. The environmental predictors associated to each cluster are displayed with alternating black and grey fonts. The purple, green and yellow circles indicate whether an environmental predictor is a major predictor of coccolithophore Shannon index patterns, computed from abundance, MAGs assemble first, or MAGs predict first method.

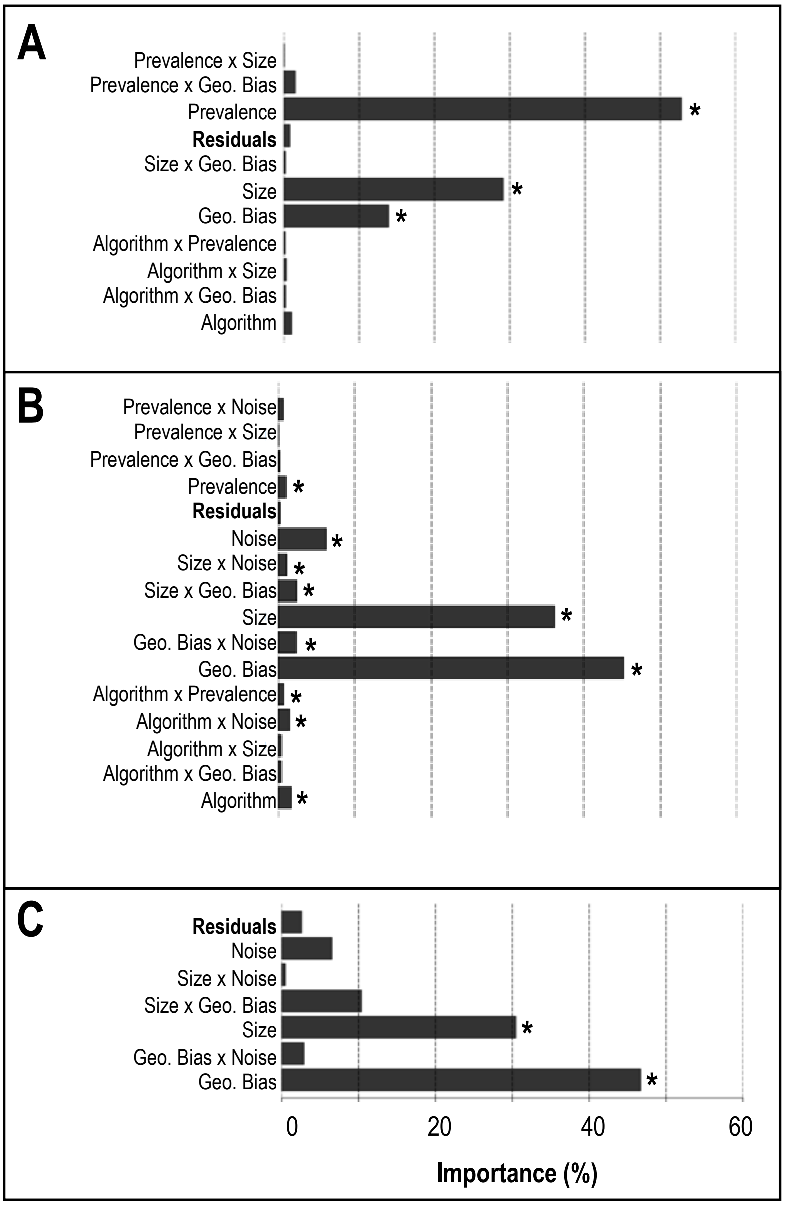


**Figure S2**: Analysis of variance of the factors influencing the true predictive performance of the habitat modelling framework for binary (**A**), continuous (**B**) and proportions (**C**) data. The asterisk indicates a significant effect (p-value < 0.05). The bars represent the importance of each effect, computed as the relative corresponding average sum of square.

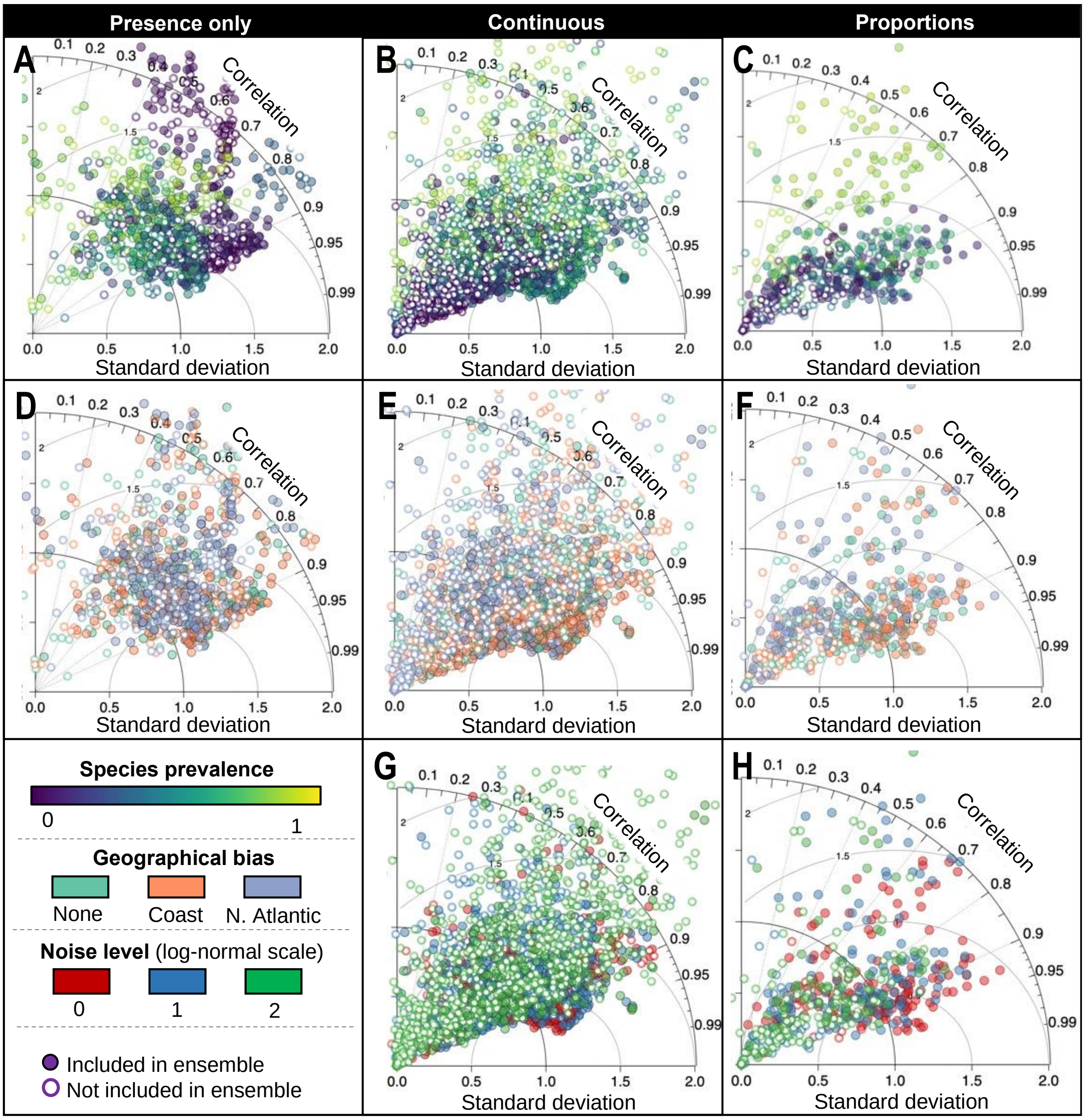


**Figure S3**: Habitat modelling performance on 20 virtual species for presence only (**A**, **D**), continuous (**B**, **E, G**) and proportion data (**C**, **F**, **H**). Each panel compare the estimated and true distribution. Each circle corresponds to a combination of prevalence, sample size, background noise, geographical bias, and algorithm. Each row is coloured according to species prevalence (**A, B, C**), geographical bias (**D, E, F**) and noise level (**G, H**). Note that presence only data cannot integrate noise bias, as all values are set to 1.

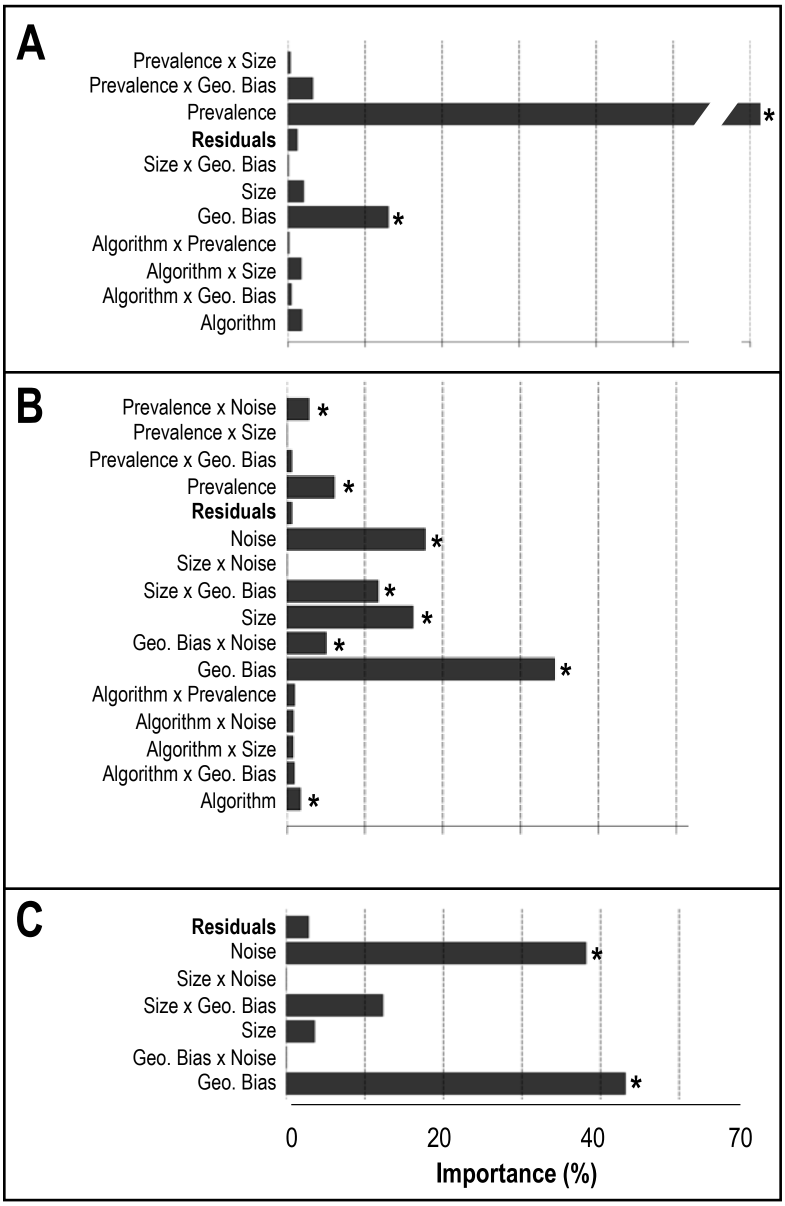


**Figure S4**: Analysis of variance of the factors influencing the false positive rate in performance estimation by the habitat modelling framework for binary (**A**), continuous (**B**) and proportions (**C**) data. The asterisk indicates a significant effect (p-value < 0.05). The bars represent the importance of each effect, computed as the relative corresponding average sum of square.

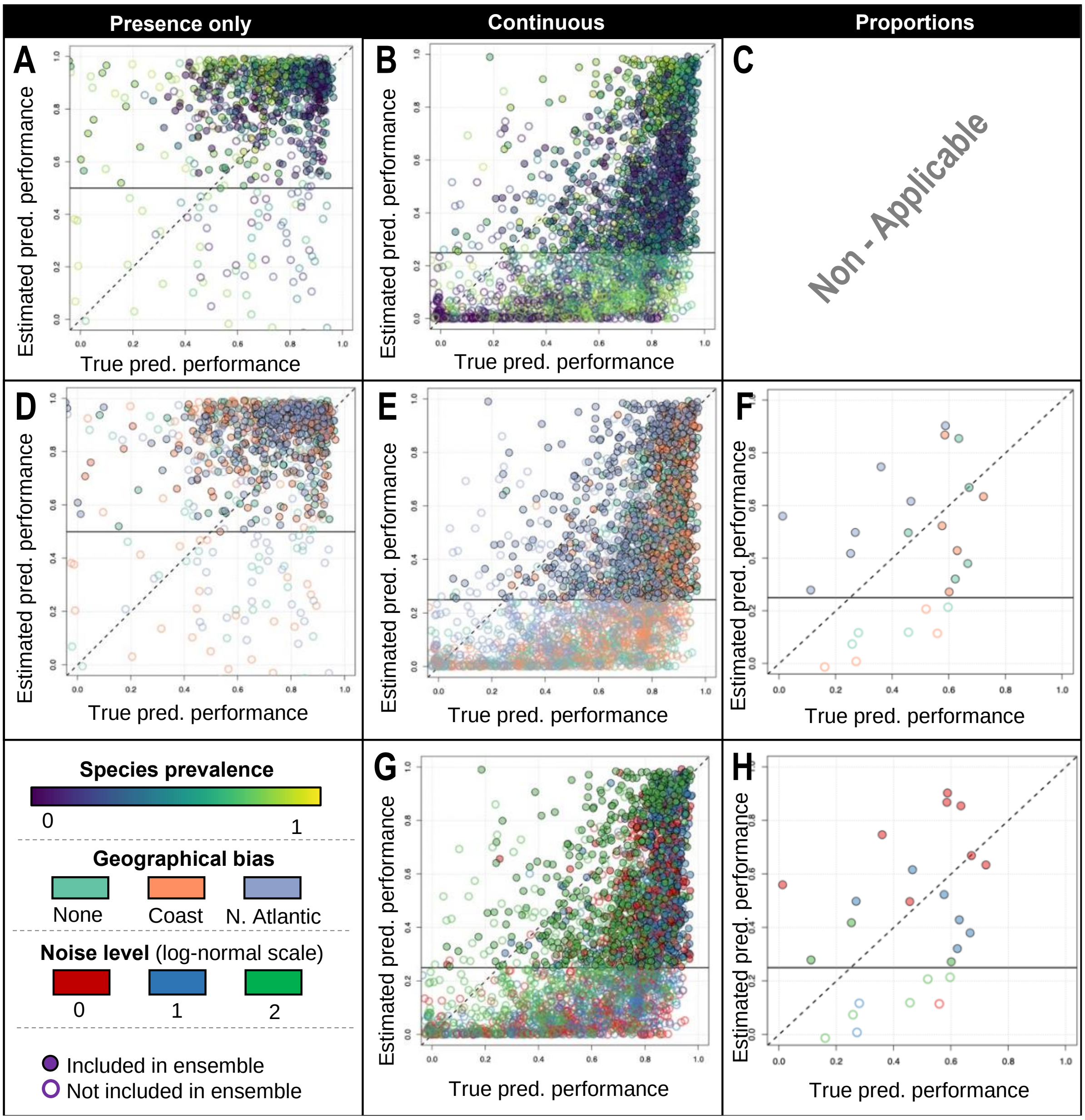


**Figure S5:** Habitat modelling performance on 20 virtual species for presence only (**A**, **D**), continuous (**B**, **E**, **G**) and proportion data (**F, H**). Each panel compare the estimated predictive performance (i.e., observed vs predicted evaluation split) to the true performance (i.e., estimated vs true distribution). The 1:1 dashed line indicates a perfect performance estimation. The horizontal black line indicates the threshold under which an algorithm is discarded.  Each circle corresponds to a combination of prevalence, sample size, background noise, geographical bias, and algorithm. Each row is coloured according to species prevalence (**A, B, C**), geographical bias (**D, E, F**) and noise level (**G, H**). Note that the estimated predictive performance is computed simultaneously across all targets (i.e., species) due to the multivariate nature of proportion data (**C**).

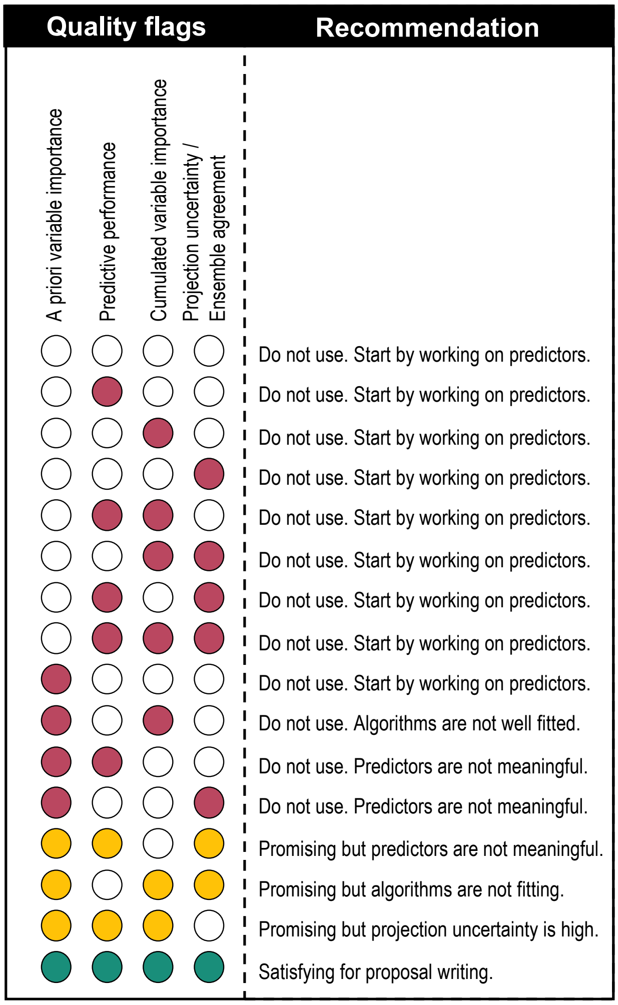


**Figure S6**: Combinations of quality flags and recommendations provided in the output of CEPHALOPOD as a traffic light system. Each circle represents one quality flag, that is coloured if successful. The red, yellow, or green colour is defined by the number of successful quality flags. A short recommendation is provided to guide the user towards an improved model.
